# Physics-Informed Ellipsoidal Coordinate Encoding Implicit Neural Representation for high-resolution volumetric wide-field microscopy

**DOI:** 10.1101/2024.10.17.618813

**Authors:** You Zhou, Chenyu Xu, Zhouyu Jin, Yanqin Chen, Bei Zheng, Meiyue Wang, Bo Xiong, Xun Cao, Ning Gu

## Abstract

Wide-field fluorescence microscopy through axial scanning provides a simple way for volumetric imaging of cellular and intracellular activities, but the optical transfer function (OTF) of wide-field microscopy suffers from axial frequency deficiencies, leading to strong interference from out-of-focus fluorescence signals and reduced imaging quality. Richardson-Lucy (RL) deconvolution and its variants are commonly employed to reduce inter-plane signal interference of wide-field microscopy. However, these methods are still affected by the “missing cone” issue inherent in the OTF, compromising both the axial resolution and optical sectioning capability. Existing deep learning methods could realize high-fidelity 3D image stack restoration, but relying on high-quality paired datasets or specific assumptions about sample distributions. Here, we propose a novel method named physics-informed ellipsoidal coordinate encoding implicit neural representation (PIECE-INR), to tackle the challenges of background signal interference and resolution loss in axial scanning image stacks using the wide-field microscopy. In PIECE-INR, we integrate the wide-field fluorescence imaging model with the self-supervised INR network for high-fidelity reconstruction of 3D fluorescence data without the need of additional ground truth data for training. We further design a novel ellipsoidal coordinate encoding based on the system’s OTF constraints and incorporate implicit priors derived from the physical model as the loss function into the reconstruction process. Our approach enables block-wise reconstruction of large-scale images by using localized physical information. We demonstrate state-of-the-art performance of our PIECE-INR method in volumetric imaging of live HeLa cells, large-volume *C. elegans* whole-embryo, and mitochondrial dynamics.

## Introduction

Fluorescence volumetric microscopy techniques^1-3^ are highly advantageous for observing cellular and intracellular activities due to their high specificity and sensitivity, as well as their ability to reconstruct the three-dimensional (3D) distribution of specific substances within a sample. Most of these methods enhance optical sectioning by restricting either the excitation or emitted light, which improves 3D imaging capabilities but at the cost of introducing complex and expensive optical systems. Wide-field fluorescence microscopy^4^, on the other hand, is simple and relatively inexpensive, also allowing for volumetric imaging of samples through axial scanning. Nevertheless, the optical transfer function (OTF) of wide-field fluorescence systems suffers from axial frequency deficiencies, known as the “missing cone” problem, leading to strong interference from out-of-focus fluorescence signals and reduced imaging quality^5^.

Deconvolution methods, such as the Richardson-Lucy (RL) algorithm^6,7^ and its variants^8-14^, are commonly employed to alleviate these effects, reducing inter-plane signal interference to a certain extent and improving the quality of 3D volumetric imaging. However, these methods, which impose physical constraints directly in iterations, are still affected by the “missing cone” issue inherent in the OTF of wide-field fluorescence systems. This leads to missing axial frequency components in the reconstructed images, thereby compromising both the axial resolution and optical sectioning capability of the results. Additionally, during the iterative process, these methods tend to amplify noise, reducing the fidelity of the solution^14,15.^ Consequently, the use of these methods often requires fine-tuning of parameters to balance the precision of the reconstruction with the noise levels, introducing significant additional workload to the process.

Recently, deep learning approaches^16,17^ have emerged, learning the mapping between input and output data across numerous samples, thus providing an automated parameter selection framework. These methods have shown remarkable capabilities in applications like super-resolution microscopy^18,19^, image denoising^20-25^, and deconvolution^26-28^, with examples including: the content-aware image restoration network (CARE) based on the U-net architecture^29^, the Richardson–Lucy Network (RLN)^26^ derived from the RL deconvolution algorithm^6,7,^ the self-Net designed for isotropic self-supervised reconstruction of samples^30^, and the residual channel attention network (RCAN)^31^. However, these methods often rely on high-quality paired datasets or specific assumptions about sample distributions, limiting their broader application. Neural fields or implicit neural representation (INR)^32^, as an emerging technology in graphics and 3D reconstruction, can be combined with physical propagation models to form a self-supervised learning paradigm that does not require large-scale additional data. INR has initially shown its potential in multiple scenarios of optical microscopic applications, such as Coordinate-based neural representations for Computational Adaptive optics (CoCoA)^33^, Diffractive Neural Field (DNF)^34^, MicroDiffusion^35^, Deep Continuous Artefact-free Refractive-index Field (DeCAF)^36^, Fourier Ptychographic Microscopy with Implicit Neural Representation (FPM-INR)^37^, Neural-Field-assisted Transport-of-intensity Phase Microscopy (NFTPM)^38^, and Semantic redundancy based Implicit Neural Compression guided with Saliency map (SINCS)^39^.

In this work, we integrate the wide-field fluorescence imaging model with the self-supervised INR network, proposing a physics-informed ellipsoidal coordinate encoding implicit neural representation (PIECE-INR) to tackle the challenges of background noise and resolution loss caused by inter-layer interference in 3D samples during wide-field fluorescence imaging. PIECE-INR establishes a self-supervised INR network for high-fidelity reconstruction of 3D fluorescence data without the need for additional ground truth data for training. We also design a novel ellipsoidal coordinate encoding method, based on the OTF constraints of wide-field microscopy systems, incorporating implicit priors derived from the physical model into the reconstruction process. This approach enables block-wise reconstruction of large-scale images using localized physical information, thereby extending the imaging field of view (FOV). We further adopt GPU parallelism to improve computing speed. Extensive experiments on both simulated and real datasets demonstrate that our method significantly enhances the reconstruction quality of wide-field fluorescence volumetric imaging, effectively mitigating the “missing cone” problem in axial scanning. Detailed comparisons with the classical RL deconvolution algorithm and the recently proposed Deconwolf method^40^ reveal that our approach substantially improves both the axial resolution and optical sectioning capabilities.

## Results

### Workflow of PIECE-INR and principle of ellipsoidal encoding

We implement PIECE-INR, a self-supervised learning method, for 3D volumetric reconstruction from multiple z-stacks captured by wide-field fluorescence microscopies (Fig. 1). The workflow of PIECE-INR consists of three main aspects (Fig. 1b). (1) Imaging process representation: PIECE-INR is designed to represent the 3D volume of samples using a coordinate-based neural network, or implicit neural representation (INR). It models the axial-scanning imaging process of multiple z-stacks as a 3D convolution by the ground-truth volume and 3D point spread function (PSF). (2) Ellipsoidal encoding: based on physics-informed constraints from the imaging process, we design a novel ellipsoidal coordinate encoding method for coordinate representation of multiple z-stack images. After self-supervised training and inferring, it achieves (near-) diffraction-limited volumetric resolution, especially with high numerical aperture (NA) objectives. (3) Boundary effect suppression: to process the large FOV imaging, we propose a method that suppresses boundary effects and enables block-wise reconstruction. This approach achieves automatic lateral and axial boundary handling with minimal artifacts. We demonstrate that PIECE-INR outperforms two most popular deconvolution tools.

**Fig. 1.**
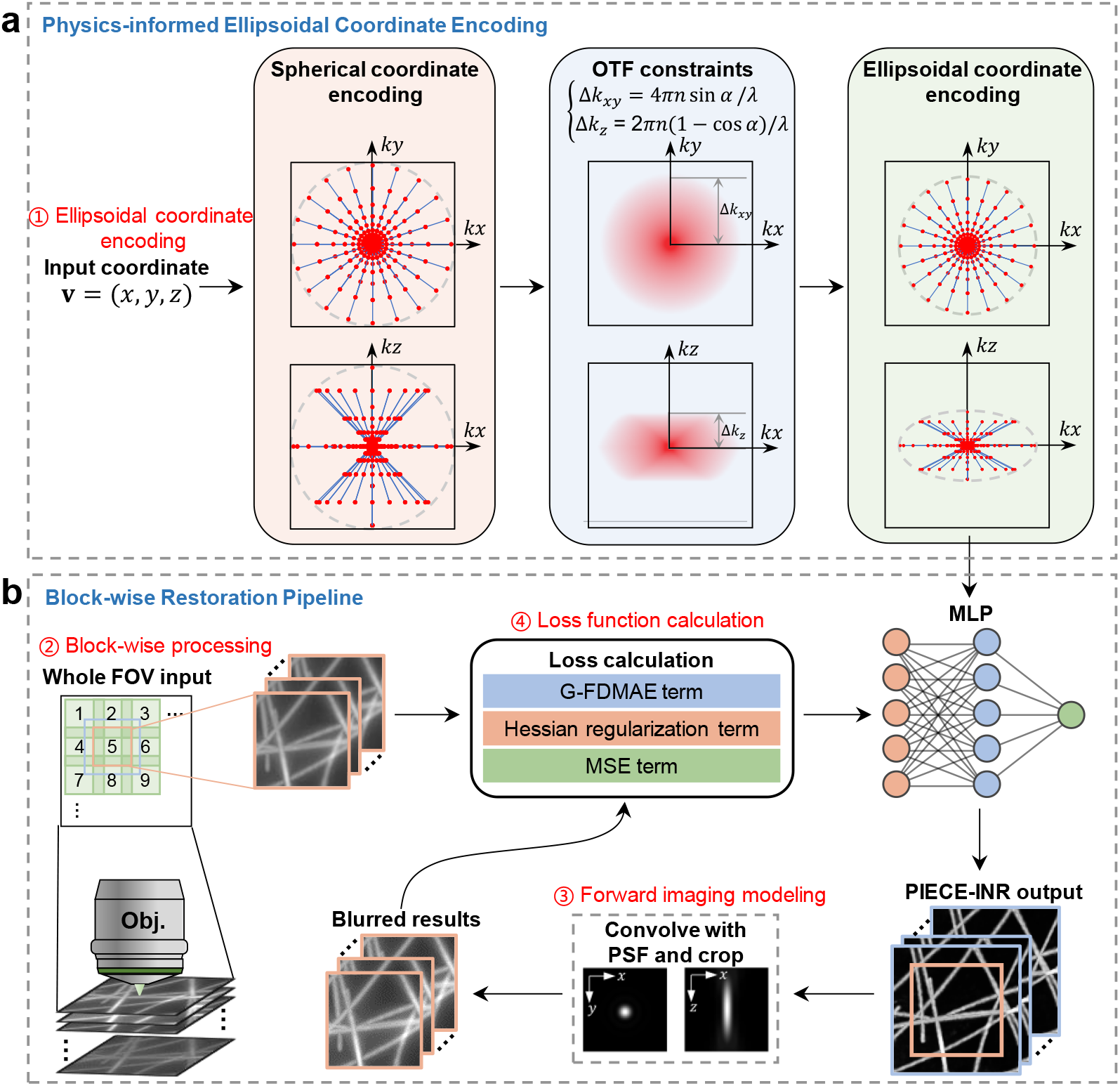
Principle of ellipsoidal encoding and pipeline of PIECE-INR. **a**, Spherical encoding ensures equal information representation in all three dimensions. Ellipsoidal encoding is used to limit the cutoff frequency of the coordinate encoding, matching the imaging physical constraints by OTF of the wide-field microscopy system. **b**, The schematic of the training pipeline of PIECE-INR, including: ➀Ellipsoid coordinate encoding, ➁block-wise processing, ➂ forward imaging modeling and ➃ loss function calculation. PIECE-INR: physics-informed ellipsoidal coordinate encoding implicit neural representation, OTF: optical transfer function, MLP: multilayer perceptron, G-FDMAE: Gaussian-Fourier domain mean absolute error.

PIECE-INR first transforms the input coordinates into encoded representations using a non-trainable ellipsoidal expansion (Fig. 1a). As shown in Fig. 1b, these encodings are subsequently mapped to the corresponding 3D structures of the sample at the input spatial coordinates through a multilayer perceptron (MLP). The output 3D structures then undergo a forward imaging process, i.e., the 3D convolution, to generate the corresponding stack of wide-field images, which are compared to the measurements to calculate the loss. The gradients are then backpropagated to update the parameters of the MLP, completing the training process. We incorporate key parameters of our microscope, including the OTF, the objective lens, voxel size, and emission wavelength λ, into the forward model. PIECE-INR learns directly from the input image stack without the need for externally labeled data and enables to reconstruct the sample’s volumetric structure with a resolution closely approximating the diffraction limit of the objective lens.

The concept of our proposed ellipsoidal coordinate encoding is visually depicted in Fig. 1a. The input consists of a set of 3D spatial coordinates, denoted as 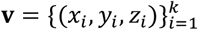, which are sampled from a pre-defined grid. Conventional encoding methods, such as Fourier positional encoding^32^ and random Fourier encoding^41^, are plagued by insufficient or anisotropic frequency support. Radial encoding^36^, while effective in capturing lateral sample features, falls short in representing axial information. In comparison, based on the assumption of isotropic nature of object in 3D space, we first propose a coordinate encoding method, termed spherical encoding, which enables isotropic information representation in all three dimensions. Furthermore, we notice that the OTF of a wide-field microscopy has different cutoff frequencies in various directions, forming an ellipsoidal shape in the frequency domain. This implies that only frequency components within the OTF’s cutoff frequency range can be effectively supported through self-supervised training process. Therefore, we further optimize and improve the spherical encoding, proposing the physics-informed ellipsoidal coordinate encoding. The novel ellipsoidal encoding can enhance the learning and representation capabilities of the network in the physical constraint (i.e., OTF-limited) effective area. This approach facilitates high-quality, artifact-free restoration of 3D sample structures, ensuring accurate and reliable results. A detailed performance comparison of different encoding methods can be found in Supplementary Fig. S3.

The purpose of PIECE-INR is to solve such an inverse problem:

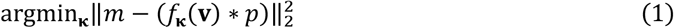

where *m* denotes the 3D measurements, *p* is the points spread function (PSF), *f* _κ_represents the MLP parameterized by weights κ, and *is the 3D convolution operator. To handle this problem, we apply the mean square error (MSE) and the Hessian regularization term, as well as our designed physics-consistency term, namely Gaussian-Fourier domain mean absolute error (G-FDMAE), as the loss function for gradient back-propagation and network parameter updates (**Methods**).

INR typically generates a significant amount of intermediate storage overhead during inference and training, which scales linearly with the size of the coordinate space^36^ and becomes further exacerbated by the introduction of ellipsoidal encoding. For 3D reconstruction of large FOV samples, block-wise processing becomes necessary. However, inter-layer optical crosstalk in multi-stack measurements causes each FOV to contain not only in-focus information but also out-of-focus light from surrounding areas. Direct block-wise processing can lead to ringing artifacts at the edges, potentially causing the entire reconstruction to collapse^42^. Traditional deconvolution methods^43^ often mitigate this effect by using constant mirroring or padding, and the Bertero-Boccacci method (BBM)^10^ addresses this issue by predefining the PSF size and adaptively predicting object information across the remaining FOV. However, applying these boundary processing methods in our PIECE-INR approach either causes the physical model to deviate too far from reality, leading to reconstruction errors, or incurs significant computational cost and memory pressure. Therefore, we propose an automatic method that efficiently suppresses boundary effects and enables block-wise reconstruction of large-volume samples (**Methods**). More details of the overall PIECE-INR approach can be found in Supplementary Note 1, Supplementary Fig. S1, and Supplementary Fig. S2.

### Evaluation and characterization of PIECE-INR

To characterize and evaluate the performance of PIECE-INR, we first simulate the wide-field microscopy image stacks of randomly 3D distributed fluorescent beads and compare the performance of PIECE-INR with traditional RL deconvolution (20 iterations) and the state-of-the-art Deconwolf method (100 iterations) using the same PSF (Fig. 2a). This simulation employs a 40x/0.75 NA air objective, featuring an axial scanning with 200 nm intervals. The specific parameters for each experiment are listed in Supplementary Table S1. We show the x-y and x-z maximum intensity projections (MIPs) of recovered beads by different methods in Fig. 1a. We find that the RL deconvolution and Deconwolf method produce results with non-uniformity in the axial direction, where the shapes of the beads at different depths are not consistently recovered. In contrast, the results of the proposed PIECE-INR maintain a high degree of consistency at different depths with reference to the ground truth, which are generated with diffraction-limited resolution, defined by the NA of the objective lens (as highlighted by yellow and red arrows in Fig. 2a). The beads restored by the Deconwolf method have lost their original spherical shape and energy distribution, which is more evident in the x-z MIP (Fig. 2a and Supplementary Fig. S4). We speculate this is due to some degree of overfitting. More intuitively, in Fig. 1b, we compare the intensity profiles of beads using different methods with the ground truth along the lines indicated by yellow dashes in Fig. 1a. In the line profile, the red curve (result of PIECE-INR) is closest to the black curve (ground truth), while the green curve (result of RL deconvolution) and the blue curve (result of Deconwolf) exhibit certain deviations from the ground truth in terms of intensity and full width at half maximum (FWHM), demonstrating PIECE-INR’s ability to uniformly represent and recover the entire 3D space, thereby avoiding potential reconstruction artifacts and errors (**Methods**).

**Fig. 2.**
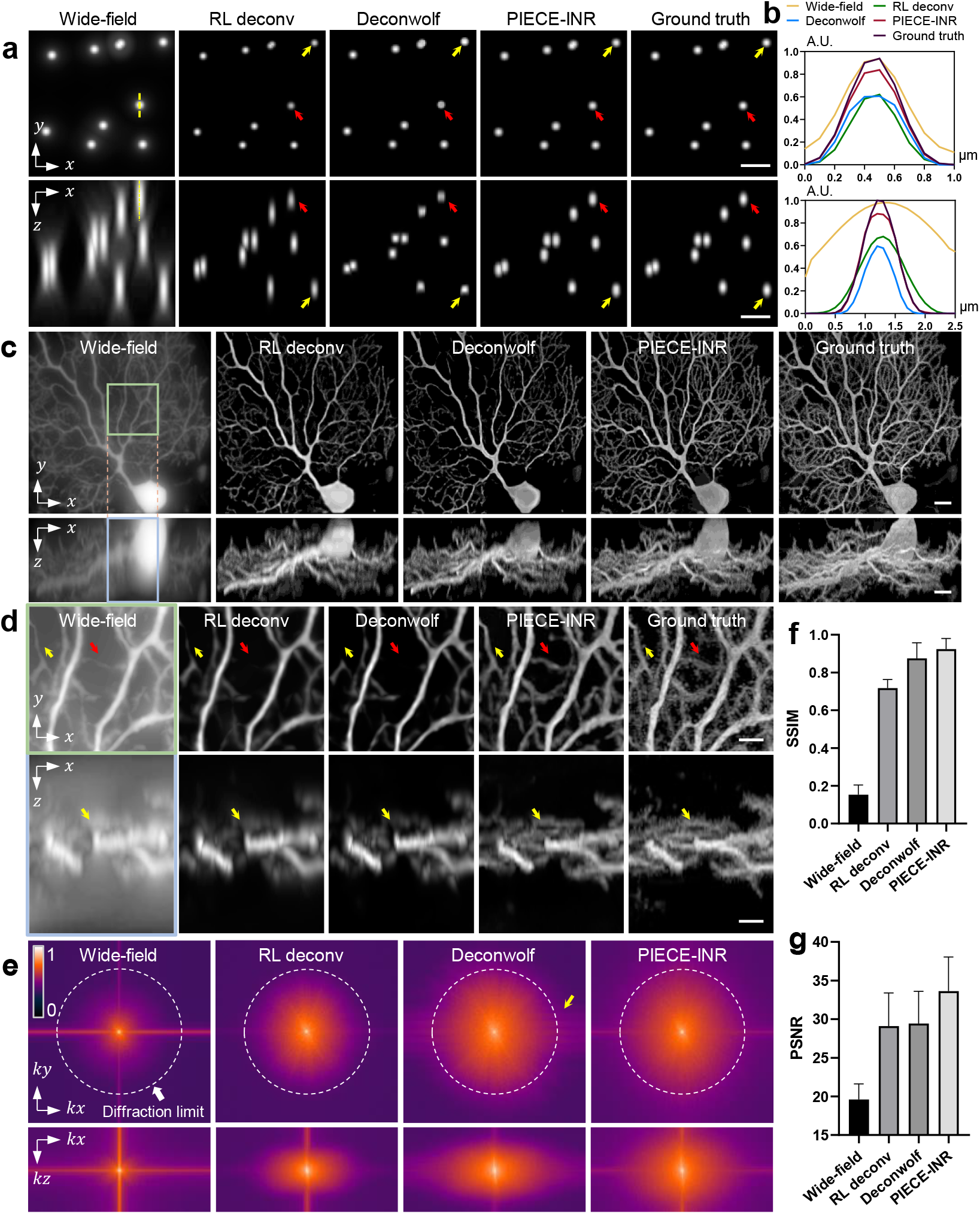
Comparison of PIECE-INR with the classical Richardson-Lucy (RL) deconvolution and the state-of-the-art Deconwolf method. **a**, x-y and x-z Maximum intensity projections (MIPs) of synthetic fluorescent beads imaged by wide-field microscopy, RL deconvolution, Deconwolf, and the proposed PIECE-INR, respectively, with the reference to the ground truth. **b**, Comparison of the intensity profiles of wide-field microscopy (yellow), RL deconvolution (green), Deconwolf (blue), and PIECE-INR (red) with the ground truth (black) along the lines indicated by the yellow dashes in **a. c**, MIPs of synthetic wide-field image stacks from Purkinje cells acquired with a confocal microscope (data source: ^44^), restored using different methods. **d**, Zoom-in areas in **c** marked by green and blue boxes, respectively. **e**, The Fourier transform of each MIP image in **c**, where the white dash lines show the diffraction limit of objective lens (40×/0.9NA). **f-g**, Peak signal-to-noise ratio (PSNR) and structural similarity index (SSIM) of the full field of view (FOV) Purkinje cells restored by different methods across the depth range, with referring to the ground truth. Source data are provided in the Source Data files. Scale bar, 2 μm (**a, d**), 5 μm (**c**).

Next, we test and characterize the performance of PIECE-INR in reconstructing high-density samples. We simulate wide-field image stack from confocal microscopy data of mouse Pukeinje cells (data source: ^44^) using a 40x/0.9 NA air objective with a 500 nm axial scanning interval. We also compare the performance of revered MIPs by applying PIECE-INR with RL deconvolution (20 iterations) and Deconwolf method (50 iterations) using the same PSF (Fig. 2c, d). We crop a small block region from the original 3D stack and show the MIP results of enlarged images in Fig. 2d. As indicated by the yellow and red arrows in Fig. 2d, PIECE-INR significantly enhances the contrast and resolution of the wide-field data and resolves the fine structures of Purkinje cell dendrites. In comparison, RL deconvolution and Deconwolf lose a substantial amount of image detail during the reconstruction process. Moreover, after transforming in to Fourier domain, PIECE-INR exhibits its ability to recover high-frequency axial information significantly surpasses that of RL deconvolution. The spatial frequency spectrum range recovered by Deconwolf result is similar to that of PIECE-INR in the lateral direction; however, it exhibits noticeable periodic artifacts (as indicated by the yellow arrows in Fig. 2e) and has a significantly narrower frequency spectrum range in the axial direction. For quantitative comparison, we calculate the peak signal-to-noise ratio (PSNR) and the structural similarity index (SSIM) of 3D volumes recovered by different methods, which also indicate that our PIECE-INR method achieves substantially better results than those resolved using RL deconvolution and Deconwolf (Fig. 2f, g). From the simulation results, we demonstrate that PIECE-INR achieves high-fidelity restoration capability while attaining lateral and axial resolutions at the diffraction limit of the objective lens, which is benefiting from the ellipsoidal encoding and customized loss function (**Methods**). PIECE-INR enables high-resolution recovery of the sample in all three dimensions, which is critical to research on complex biological structures^45^, cellular dynamics^46-48^, and tissue organization^49^, where subtle details and precise spatial relationships are crucial for understanding underlying mechanisms and behaviors.

### PIECE-INR accomplishes high-fidelity and artifact-free 3D restoration

Fidelity is a key metric for evaluating computational microscopy algorithms^50^, but maintaining it in 3D wide-field image restoration is challenging. This is due to two main factors: the significant crosstalk between different image layers, and the reliance of deconvolution methods like RL and Deconwolf on Poisson noise models and maximum likelihood estimation (MLE). Given that noise is inevitably introduced in images captured during practical experiments, excessive iterations often lead to solutions dominated by noise^14^, causing detail loss and noise amplification (Supplementary Fig. S5). Enhanced by incorporating prior knowledge, the proposed ellipsoidal encoding and customized loss function (**Methods**) enable near-diffraction-limit resolution and high-fidelity restoration.

To verify the performance of PIECE-INR, we first image a live HeLa cell co-expressing Tub-GFP and Tom20-mCherry using a 100×/1.5NA oil-immersion objective with a 200 nm axial scanning interval (Fig. 3a-d). As illustrated in the x-y and x-z MIP images (Fig. 3a), PIECE-INR significantly enhances the signal-to-background ratio (SBR), contrast, and resolution of images captured by the wide-field microscopy. From the enlarged images shown in Fig. 3b and 3c, Deconwolf (100 iterations) tends to amplify noise when solving dense cellular structures, resulting in a grainy appearance, while the results of PIECE-INR maintain good continuity with fine details and no shot noise. This observation is further supported by Fig. 3d, which compares the x-z direction slices of the green channel (microtubule) and the cherry channel (mitochondria) using Deconwolf (top) and PIECE-INR (bottom), as indicated by the yellow dashed lines in Fig. 3a. The Deconwolf result is substantially corrupted by heavy background, obscuring the sample, while the PIECE-INR restoration shows a clean background and well-defined sample structure.

**Fig. 3.**
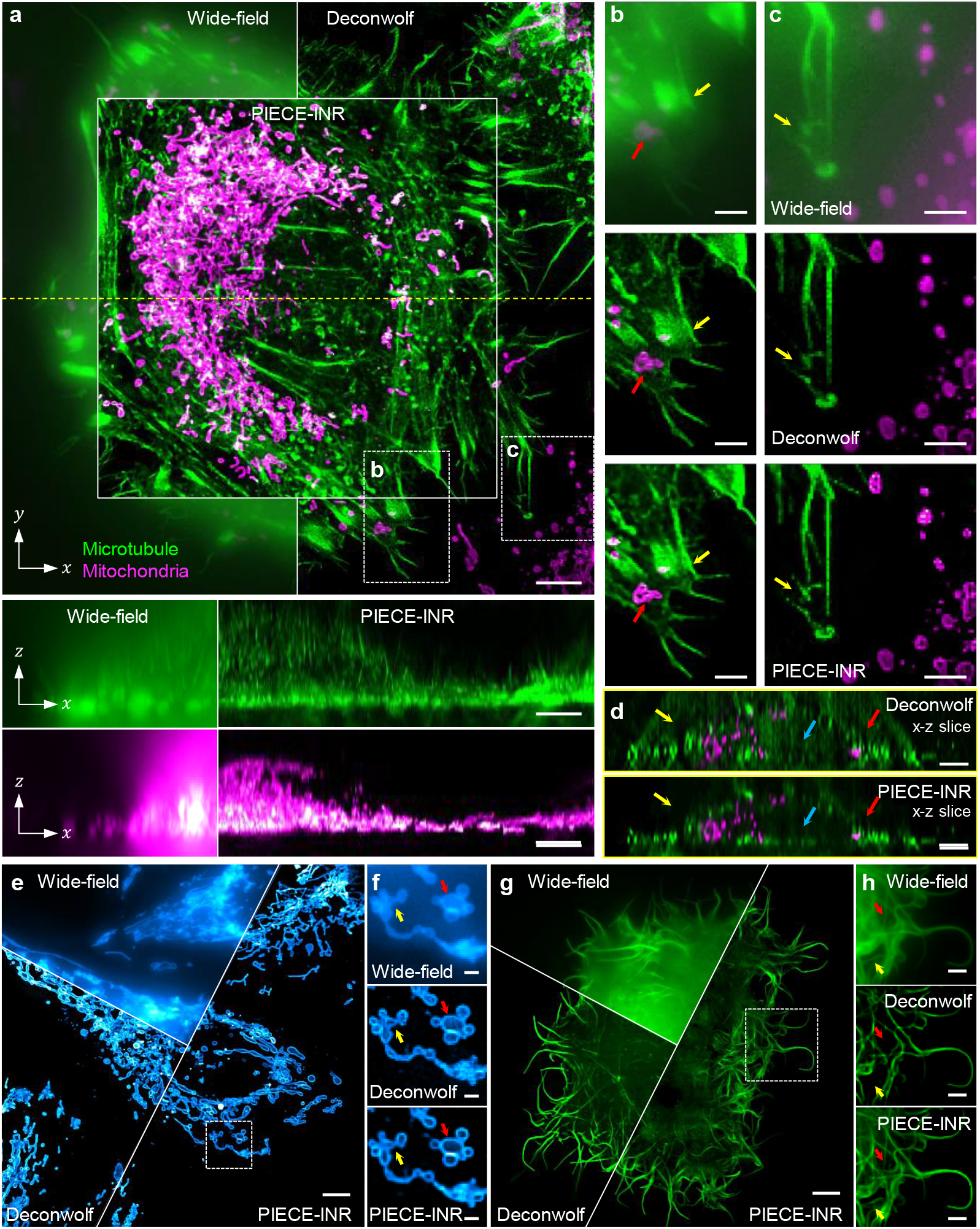
Experimental verification of PIECE-INR across various live samples. **a**, Representative restored images of the microtubule cytoskeleton and mitochondria in a live HeLa cell (100×/1.5NA oil-immersion objective) co-expressing Tub-GFP and Tom20-mCherry, processed using Deconwolf and PIECE-INR. Both x-y and x-z MIPs are shown for comparison of raw wide-field images and images processed by Deconwolf and PIECE-INR. **b-c**, Zoom-in areas in **a** marked by white boxes. **d**, Comparison of x-z direction slices of the green channel (microtubule) and the cherry channel (mitochondria) using Deconwolf (top) and PIECE-INR (bottom), as indicated by the yellow dashed lines in **a. e**, Representative restored images of mitochondria in a live HeLa cell expressing mCherry-mito (100×/1.5NA oil-immersion objective). **f**, Zoom-in areas in **e** marked by white boxes. **g**, Representative restored images of the microtubule cytoskeleton in a live COS-7 cell expressing Tub-GFP (100×/1.5NA oil-immersion objective). **h**, Zoom-in areas in **g** marked by white boxes. Scale bar, 1 μm (**h**), 2 μm (**b, c, f**), 5 μm (**a, d, e**).

We also observe the superior performance of PIECE-INR in imaging live HeLa cells labeled with mitochondrial markers (Fig. 3e, f) and live COS-7 cells labeled with microtubule markers (Fig. 3g, h), with data are captured using the same 100×/1.5NA oil-immersion objective with a 200 nm axial scanning interval. PIECE-INR exhibits greater continuity and reduced shot noise when imaging fine structures, as highlighted by the red and yellow arrows in Fig. 3f. Furthermore, Deconwolf often over-sharpens sparse linear structures, leading to excessive refinement or even loss of original sample details. PIECE-INR, however, preserves the original details appeared in the wide-field images while enhancing overall image quality (Fig. 3c, g-h). In summary, PIECE-INR demonstrates high reliability and stability, converging to the optimal solution as training progresses. PIECE-INR also imposes no restrictions on observed samples and consistently delivers exceptional restoration results, whether dealing with dense structures or sparse point-like or linear features.

### PIECE-INR well resolves the high-density, large-volume *C. elegans* whole-embryo

To further test PIECE-INR’s capability in resolving dense samples, we employ an open-source dataset of *C. elegans* whole-embryo (data source: ^43^), which presents a challenging scenario due to its complex structure, large volume, and multiple color channels. The fluorescence image dataset comprises three color channels (DAPI, FITC, and CY3), each labeling distinct cellular structure: chromosomes in the nuclei, microtubules, and protein spots, respectively. As shown in Fig. 4a, the images restored by PIECE-INR have much higher structural richness and higher contrast in comparison with RL deconvolution (20 iterations) and Deconwolf (50 iterations). Notably, our approach to mitigating boundary effects (**Methods**) allows for region-by-region data processing, enabling boundary-effect-free restoration over the whole volume, as shown in the x-y MIP images (Fig. 4a, b). We also conduct a comparative analysis of our method against cases without boundary suppression and other boundary handling techniques (Supplementary Fig. S8).

**Fig. 4.**
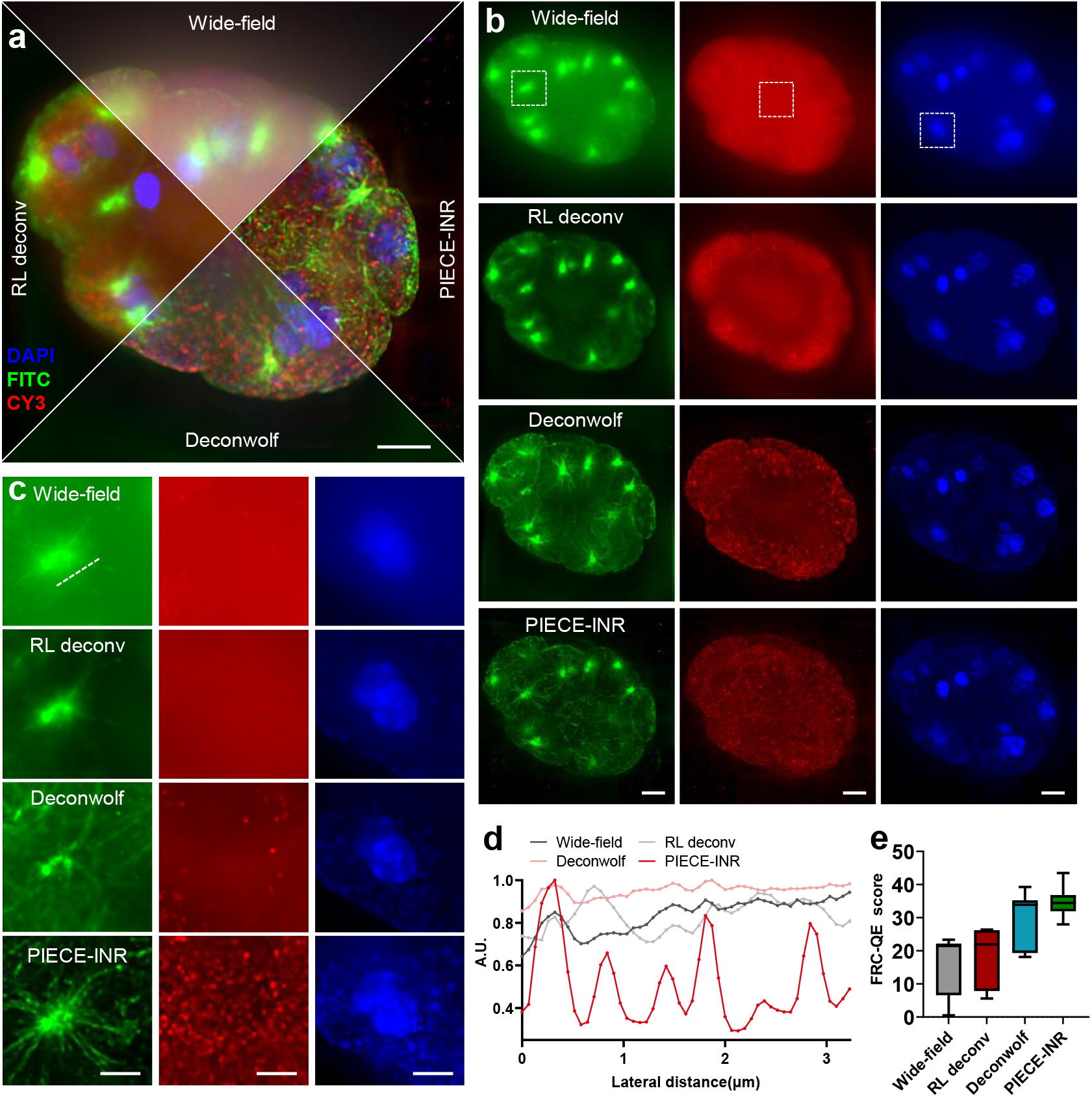
PIECE-INR greatly enhances 3D restoration quality of high-density, large-volume *C. elegans* whole-embryo image stacks. **a**, Multi-color MIP images recovered using RL deconvolution, Deconwolf, and PIECE-INR respectively, compared to raw wide-field images (100×/1.4NA oil-immersion objective). The *C. elegans* whole-embryo, with a 43.3×43.3×8μm^3^ volume, is simultaneously labeled with DAPI (461 nm), FITC (519 nm), and CY3 (570 nm). **b**, Full-FOV MIP images of three channels restored using different methods, compared to the wide-field images. **c**, Zoom-in areas marked by white boxes in **b. d**, Comparison of intensity profiles for wide-field (black), RL deconvolution (gray), Deconwolf (pink), and PIECE-INR (red) along the white dashed lines in **c. e**, Statistical comparison of wide-field, RL deconvolution, Deconwolf, and PIECE-INR in terms of resolution, measured by the Fourier ring correlation quality estimate (FRC-QE) score in the FITC channel. The plot displays medians (center line), 25th and 75th percentiles (box limits), and maximum and minimum values (whiskers). Source data are provided in the Source Data file. Scale bar, 2 μm (**b, c**), 5 μm (**a**).

PIECE-INR can effectively recover complex structural information of samples from wide-field images with severe inter-layer crosstalk. As shown in the FITC-labeled fluorescence images (left panel of Fig. 4c), a microtubule aster structure is visible in the wide-field image of the *C. elegans* embryo, albeit with very poor contrast. PIECE-INR clearly resolves the microtubule aster structure with high contrast, fine details, and a clean background, while RL deconvolution and Deconwolf fail to recover this structure. The intensity profiles along the white dashed lines in Fig. 4c further support this conclusion, as PIECE-INR clearly distinguishes the microtubules radiating from the microtubule aster, whereas the other methods do not (Fig. 4d). In the middle panel of Fig. 4c, the intensity of CY3-labeled protein spots is weaker than the background intensity in the wide-field MIP image, causing structural information to be overwhelmed by the out-of-focus signal in the background. This severe inter-layer crosstalk prevents RL deconvolution from recovering the original structure in this region. While Deconwolf performs better, it still fails to resolve all protein spots. In contrast, PIECE-INR clearly restores a significantly greater number of protein spots in the *C. elegans* embryo. The ability to effectively recover signals amidst significant background noise is also demonstrated by the MIP images of DAPI-labeled chromosomes in the left panel of Fig. 4c. We also employ the Fourier ring correlation quality estimate (FRC-QE)^51^ to quantitatively evaluate the spatial resolution in the FITC channel. The resolution achieved by PIECE-INR surpasses that of RL deconvolution and Deconwolf, showing greater consistency across layers with significantly lower variance (Fig. 4e). These results highlight PIECE-INR’s exceptional performance in restoring high-density, large-volume complex samples and its ability to recover structures obscured by background noise, demonstrating its robust and powerful 3D restoration capability.

### PIECE-INR enables high-precision dynamic 3D imaging of live samples using wide-field microscopy

Mitochondria are highly dynamic organelles^52^, continually undergoing two phases of fusion (where two mitochondria merge into one) and fission (where a single mitochondrion splits into two), collectively referred to as “mitochondrial dynamics”. We demonstrate here that PIECE-INR can improve the capability boundary of wide-field microscopy and achieve high-quality volumetric imaging of mitochondrial dynamics. We conduct live-cell imaging of HeLa cells expressing Tom20-mCherry using a wide-field fluorescence microscope equipped with a 100×/1.5NA oil-immersion objective, with depth information reflected through pseudo-color encoding (Fig. 5a, b). We axially scan an image stack of sample volume with 200 nm intervals, with each capture process taking 3 seconds.

**Fig. 5.**
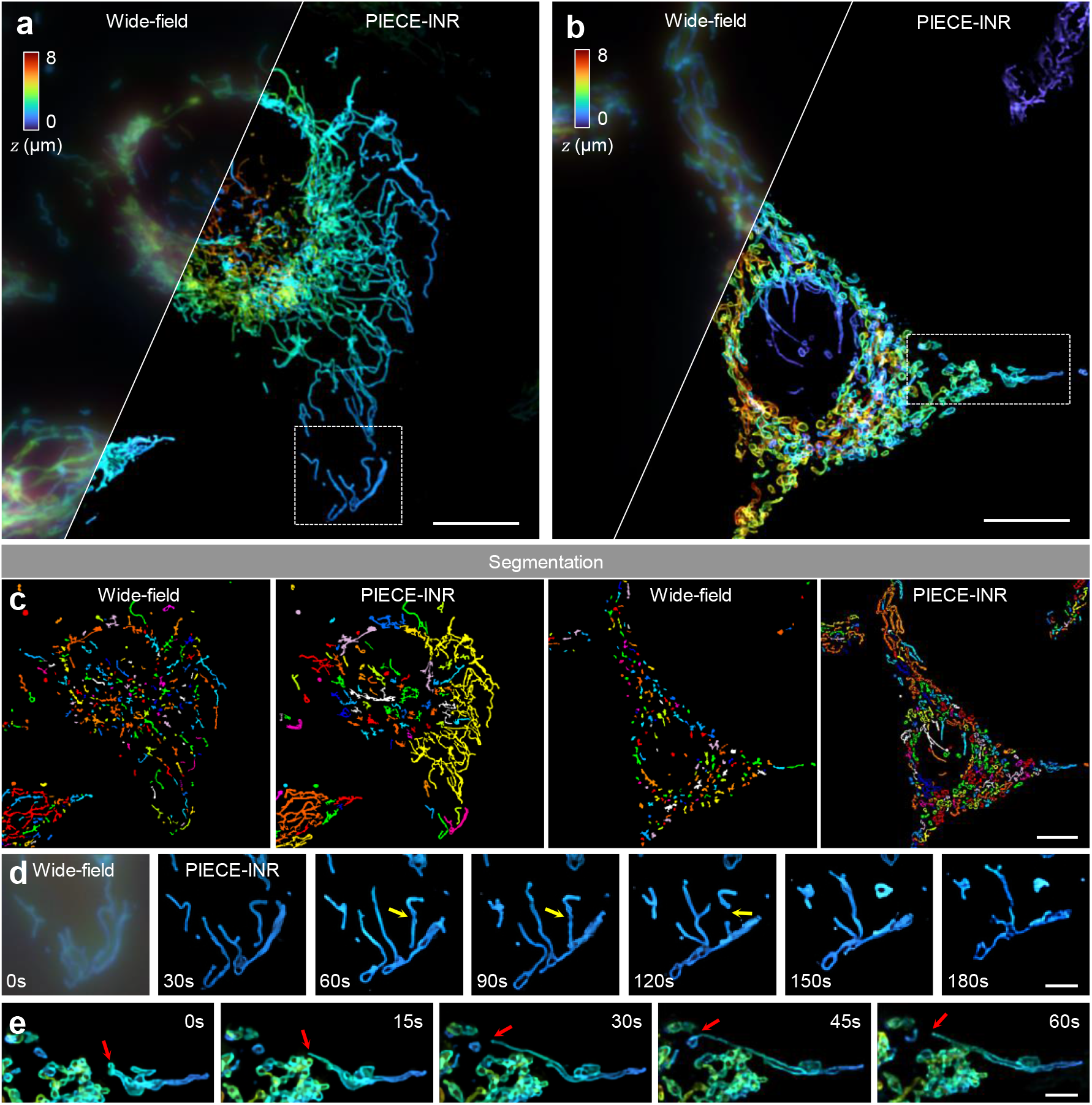
PIECE-INR enables high-resolution dynamic observation of mitochondrial life activities. **a-b**, MIP images recovered using PIECE-INR, compared to raw images acquired by a wide-field microscope of live HeLa cells expressing Tom20-mCherry (100×/1.5NA oil-immersion objective). Depth information is reflected through pseudo-color encoding. **c**, Segmentation results of the mitochondria based on the input of x-y MIPs in **a** and **b** correspondingly. **d**, Time-lapse images illustrate the dynamic process of mitochondrial fission, as indicated by yellow arrows. **e**, Time-lapse images illustrate the dynamic process of kiss-and-run events, as indicated by red arrows. Scale bar, 10 μm (**a**-**c**), 3 μm (**d, e**).

The raw wide-field images exhibit low contrast, making mitochondrial structures difficult to discern. In contrast, the PIECE-INR method effectively restores high-fidelity, high-resolution details of mitochondrial morphology. As shown in Fig. 5a, PIECE-INR effectively resolves the elongated, rod-like shape of mitochondria. While in Fig. 5b, PIECE-INR accurately reveal the toroidal cross-sections of the outer mitochondrial membrane (OMM), which are indiscernible in the raw wide-field images. By employ an open-source tool^53^, we demonstrate that compared to raw wide-field images, the resulting high-quality images empowered by PIECE-INR can enable precise segmentation (Fig. 5c). PIECE-INR also clearly illustrates the dynamic mitochondrial processes, such as fission (Fig. 5d) and kiss-and-run events^51,54^ (Fig. 5e).

## Discussion

In this work, we propose PIECE-INR, a physics-informed ellipsoidal coordinate encoding implicit neural representation, designed to address the challenges of heavy background and resolution loss in wide-field fluorescence microscopy with axial scanning. By integrating the wide-field imaging model with a self-supervised INR network, we establish a framework for high-fidelity 3D fluorescence reconstruction without requiring ground truth data. We design a novel ellipsoidal coordinate encoding based on the system’s OTF constraints and incorporate implicit priors as the loss function, enabling block-wise reconstruction of large-scale images using localized physical information. We demonstrate PIECE-INR’s effectiveness in volumetric imaging of live HeLa cells, large-volume *C. elegans* embryos, and mitochondrial dynamics. PIECE-INR holds significant potential for bioimaging applications, such as enhancing the sensitivity and spatial resolution of imaging-based spatial omics. On a technical level, further improvements will focus on increasing the algorithm’s noise robustness to reduce photobleaching, optimizing the computational architecture to accelerate large-field 3D volume reconstruction for real-time observation, and expanding its application to other live 3D imaging modalities, such as light-sheet microscopy and structured illumination microscopy.

## Methods

### Wide-field fluorescence microscope

The wide-field microscope system was constructed using an inverted fluorescence microscope (MF53-NH, MShot), which includes a four-channel fluorescent LED light source (MG120, MShot) with wavelengths of 365 nm, 460 nm, 560 nm, and 625 nm. To achieve precise axial displacement, a piezo objective scanner (Coretomorrow, P73.Z500S) was utilized. The objectives used during image acquisition included two commercial objectives: 40X/0.75NA air-immersion (UPLFLN40X, Olympus) and 100X/1.5NA oil-immersion (UPLAPO100XOHR, Olympus), both mounted on the piezo objective scanner. Images were captured by an sCMOS camera with 2048×2048 pixels and a pixel size of 6.5 μm (MSH12, MShot).

### Computational details

PIECE-INR is trained on a PC with an Intel Core i7-13700K processor and RTX 3090 graphic cards (NVIDIA) under the software environment of Pytorch2.1.1+cu118 and Python 3.9.0. When solving samples with large FOV, we use the block-wise strategy and reconstruct blocks by adopting GPU parallelism to improve computing speed. For example, for a sample with volume size of 500×500×30 pixels, we literally split it into 6×6 blocks, with each size of 100×100×30 pixels and literal margin of 20 pixels, and then assign the PIECE-INR method for each block. The sequential computation takes about 5×25 minutes to complete the whole reconstruction. By implementing parallel computing across four GPUs, we reduce the computation time to approximately 30 minutes.

### Cell culture, transfection, and staining

COS-7 (CRL-1651) cell line and HeLa cell line were purchased from the American Type Culture Collection (ATCC). The cells were cultured in DMEM (Gibco, catalog no. C11965500CP) and maintained at 37 °C with 5% CO_2_ and routinely tested for mycoplasma contamination. On the day before transfection, cells were seeded on high-precision coverslips (Marienfeld, catalog no. 0117650) that were coated with poly-L-lysine (Sigma-Aldrich, catalog no. P4707) in six-well plates. For transient transfections, the cells were transfected with Lifeact-mGreenLantern using Lipofectamine™ 3000 (Thermo Fisher Scientific, catalog no. L3000015), following the manufacturer’s instructions. For single transfections, 1 μg of DNA was used per well in a six-well plate. In the case of co-transfections with HeLa cells, 0.75 μg of each DNA sample was used, totaling 1.5 μg per well. To label actin in live cells, Lifeact-mGreenLantern was constructed and transfected into COS-7 cells. The Lifeact and mGreenLantern genes were amplified by PCR from the plasmids Lifeact-RFP (a gift from Liangyi Chen, Peking University, Beijing, China) and Clathrin-mGreenLantern (Addgene, no.164462), respectively. The mGreenLantern gene was fused to the C-terminus of Lifeact. To label mitochondria, Tom20-mCherry (a gift from Liangyi Chen, Peking University, Beijing, China) was transfected into HeLa cells. To simultaneously label actin and mitochondria in live cells, Lifeact-mGreenLantern and Tom20-mCherry were co-transfected into HeLa cells. For live cell imaging, 35 mm glass-bottom dishes (FCFC020, Beyotime) were pre-coated with 50 µg/ml collagen. Subsequently, 2×10^4^ cells were seeded onto each coverslip.

The human umbilical cord mesenchymal stem cells (hUC-MSCs) were obtained from the Clinical Stem Cell Center of the Nanjing Drum Tower Hospital in China. The cells were seeded in a 35 mm glass bottom dish with a 20 mm bottom well (Cellvis, catalog no. D35-20-1-N) and cultured under standard conditions (37 °C, 5% CO_2_). The basal growth medium (500 mL) contained 75 ml fetal bovine serum (Corning, catalog no. 35-081-CV), 5 ml antibiotics (penicillin and streptomycin) (Thermo, catalog no. 15140122), and 420 ml low-glucose DMEM (Thermo, catalog no. 10567014). For staining actin filaments (F-actin), hUC-MSCs were fixed with 4% formaldehyde solution in PBS at room temperature for 30 minutes and then washed with PBS three times. Next, 0.1% Triton X-100 in PBS was added into the fixed cells for 5 minutes to increase permeability and the permeabilized cells were washed three times in PBS. Subsequently, 1 ml 1×Phalloidin conjugate working solution (Phalloidin-iFluor 488 Reagent, abcam, catalog no. ab176753) was added into the dish and incubated for 1h at room temperature. Finally, the cells were sealed by antifade mounting medium (with DAPI) (Servicebio, catalog no. G1407).

### Spherical encoding and physic-informed ellipsoidal encoding

The spherical encoding and physics-informed ellipsoidal encoding proposed can be viewed as extensions of existing encoding methods, including Fourier positional encoding, random Fourier encoding, and radial encoding. The formula for spherical encoding is as follows:

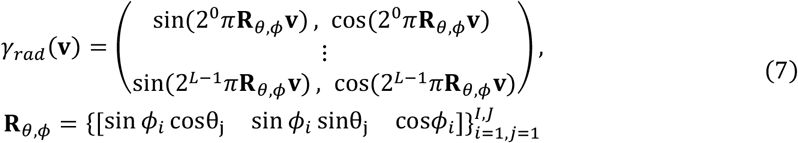

where **v** = (*x, y, z*) is the input spatial position, **R**_*θ,ϕ*_ represents the radial vector of the unit sphere, and *L* > 0denotes the total number of frequencies. *ϕ*_*i*_ and θ_*j*_ are angular parameters that determine the direction of the radial vector.

Spherical encoding provides MLP with a series of supports in the frequency domain. In addition, considering the system’s OTF, only the frequency components within the OTF cutoff frequencies *f*_*cr*_ and *f* _*cz*_ can obtain effective support in a self-supervised manner. Specifically, the cutoff frequency in the *k*_*z*_ direction is typically smaller than in the *k*_*x*_ and *k*_*y*_ directions (Fig. 1a). As a result, the OTF in the *k*_*x*_-*k*_*z*_ plane is not circular, but rather elliptical. Different physical parameter settings result in vastly different frequency domain ranges for the system’s OTF. Directly adjusting the angles and encoding depth of the spherical encoding to accommodate different physical parameters would incur a significant hyperparameter search cost. Therefore, based on spherical encoding, we design an adaptive ellipsoidal encoding method tailored to the OTF range of the wide-field microscopic system. The formula for physics-informed ellipsoidal encoding is as follows:

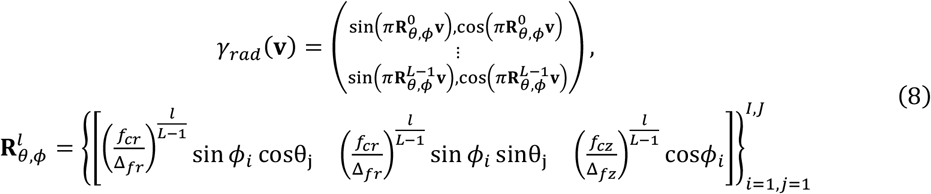

where Δ_*fr*_ and Δ*f*_*z*_ represents the frequency domain resolution corresponding to the current coordinate density. Here, **R**_*θ,ϕ*_ denotes a collection of the radial vector of the unit ellipsoid with a major axis length of 1. By varying the values of *ϕ*_*i*_ and θ_*j*_, our proposed physic-informed ellipsoidal encoding can achieve frequency extension throughout the entire coordinate space represented by the ellipsoid, effectively computing the 3D structure of a sample and its dependencies in 3D space within the diffraction limit of the objective lens.

### Mechanism and loss function of PIECE-INR

The purpose of PIECE-INR is to reconstruct the 3D structure of a sample from an image stack obtained via axial scanning with a wide-field fluorescence microscope, using self-supervised training to address the problem defined by Eq. 1. A wide-field fluorescence microscope can be modeled as a spatially invariant linear system, where the forward imaging process can be written as *m*(*x*) = *o*(*x*) *p(*x*). Here, *m* represents the image formed by the wide-field microscope, *o* is the spatial distribution of the sample’s fluorescence intensity, and *p* is the system’s spatially invariant 3D PSF. The symbol *indicates 3D convolution. Consequently, we incorporate the system’s forward imaging process into the training process to enable self-supervised training. A detailed description of the imaging model is provided in Supplementary Note 1.

We train the INR network by minimizing the error between the predicted outcomes, obtained by propagating the network’s reconstructed result through a physical process, and the actual measurements. The loss function used to compute this error is:

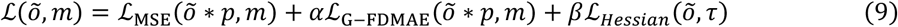

where ℒ_MSE_, ℒ_G−FDMAE_, and ℒ_*Hessian*_ represent the MSE term, the Gaussian-Fourier domain mean absolute error (G-FDMAE) term (detailed in the following paragraph), and the Hessian regularization term, respectively. Õ = *f*_θ_*(m)* denotes the predicted output of the MLP and τ is the parameter that controls the balance between lateral and axial smoothness priors. *α* and *β* are the weights associated with the ℒ_G−FDMAE_ term and ℒ_*Hessian*_ term, respectively. The MSE term is expressed as follows:

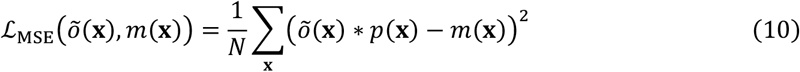

where **x** ∈ (*x, y, z*) is the pixel index of the volume in the spatial domain.

However, according to the Parseval theorem, the ℒ_MSE_ term in the spatial domain is equivalent to an MSE loss in the Fourier domain, implying that all frequency components are considered. This equivalence may lead to a loss of high-frequency details, given that both õ \**p* and *m* are theoretically band-limited signals subject to the OTF, with no effective frequency components outside the OTF’s support domain. To address this, we introduce the G-FDMAE, a modification of the FDMAE^55^ that accounts for the band-limited nature of the measurements. The G-FDMAE term is defined as follows:

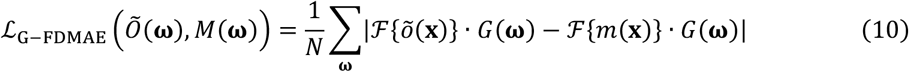

where **ω** ∈ (*k*_*x*_, *k*_*y*_, *k*_*z*_) is the pixel index of the volume in the frequency domain and ℱ{·} denotes the 3D Fourier transform. *G*(**ω**) represents a Gaussian window function, with its effective range constrained to the vicinity of the OTF’s support domain. Experimental results indicate that the inclusion of the G-FDMAE term enhances the reconstruction resolution, allowing for the recovery of finer details of the objects compared to the absence of G-FDMAE term (Supplementary Fig. S6).

The Hessian term is employed to preserve the inherent continuity of biological samples and can be expressed as follows:

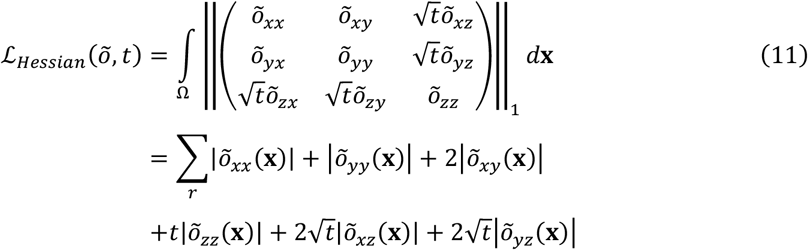

where Ω ∈ (*x, y, z*) covers all pixels of õ, *t* controls the balance between lateral and axial smoothness priors, and ‖·‖_1_ represents the 1^st^ entry-wise matrix norm.

### Block-wise training and large field-of-view reconstruction strategy

While INR typically requires small storage for the data itself, it generates substantial intermediate storage overhead during inference and training, which scales linearly with the size of the coordinate space^36^. This issue becomes further exacerbated by the introduction of ellipsoidal encoding. Furthermore, the measurements of the image stack within a single FOV contain not only information of this current region but also out-of-focus light from surrounding regions. During deconvolution, this unknown energy contribution can lead to ringing artifacts at the edges, potentially causing the entire reconstruction to collapse^42^. Traditional deconvolution methods^56^ often mitigate this effect through mirroring or padding with constant values. The Bertero-Boccacci method (BBM)^10^ addresses this issue by predefining the PSF size and adaptively predicting object information across the remaining FOV during deconvolution, thereby compensating for the impact of external signals. However, applying these boundary processing methods in our PIECE-INR approach either causes the physical model to deviate significantly from reality, leading to reconstruction errors, or imposes substantial computational and memory overhead. Therefore, we propose an automatic method that efficiently suppresses boundary effects and enables block-wise reconstruction of large-volume samples. This method includes: (1) block-wise reconstruction, (2) information-based block selection for reconstruction, and (3) suppression of boundary effects in the reconstruction process.

In the following, we begin by discussing the commonly used divide-and-conquer reconstruction for large-scale image processing, followed by the introduction of our proposed information-based block selection method. Building on this, we finally present our proposed boundary effect mitigation approach.

#### Divide-and-conquer reconstruction for large-scale samples

For a large-scale image stack, we first divide images into *m* × *n* small blocks of the same size, with some overlap between adjacent blocks (Supplementary Fig. S2). Then, we reconstruct each block individually using a separate network, utilizing only the measurements corresponding to that block and without introducing any additional information. After reconstruction, we tile the results of each block together to obtain the complete reconstruction of the large-scale sample. The edges of the tiled image are blended using a linear smoothing scheme.

#### Efficient reconstruction through information-based block selection

Given that microscopic images of biological samples, especially fluorescence signals, are often sparsely distributed and do not occupy the entire FOV, we propose a straightforward method to assess whether each block contains valid information based on its mean and variance. The specific criterion is defined by the following equation:

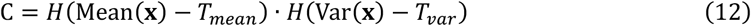

where *C* is a binary variable indicating whether a block contains sufficient information (1 if true, 0 if false). *H*(·) represents the Heaviside step function, while Mean(·) and Var(·) denote the pixel mean and variance of the block, respectively. *T*_*mean*_ and *T*_*var*_ represent the threshold values for mean and variance, respectively. Blocks that do not meet the criteria can be skipped or processed with reduced computational effort, thus reducing the overall reconstruction time (Supplementary Fig. S2). Furthermore, although our method is currently demonstrated for reconstructing samples with a complete rectangular FOV, the reconstruction process for each block is independent and decoupled, allowing for potential extension to handle measurements with irregular edges.

#### Localized physical information approximation for mitigating boundary effects

Leveraging the powerful representational capabilities of INR, we propose a localized physical information approximation method to mitigate boundary effects. This approach can effectively compensate boundary effects by predicting only a small region beyond the FOV, avoiding the need for additional predictions of large surrounding areas equivalent to the PSF size, as required in methods like BBM and Deconwolf. By experimental tests in Supplementary Fig. S7, we find that a padding of 20 pixels is sufficient to achieve stable reconstruction results. The details of our boundary effect mitigation approach are as follows:

1. Divide the wide-field measurement data, with dimensions of *D* × *W* × *H*, into smaller blocks. Uniformly partition the data into *M* × *N* overlapping blocks, denoted as *m*_*i,j*_, each with dimensions of *D* × *F* × *F*. Ensure that adjacent blocks overlap by a length of *L*, where *F* is typically set to 100 and *L* ranges from 10 to 20.
2. Assign a separate neural network 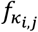 to each small block. Also, allocate a predefined coordinate grid *x* to each block, which will be used for computation. The size of the coordinate grid is set to (*D + R*) × (*F + R*) × (*F + R*), where *R* represents a small strip region outside the FOV, introduced to achieve localization of physical information, and is typically set to 20.
3. For each small block, calculate its mean and variance. Then, compute the decision value according to Eq. 12. If C < 0, it indicates that there is no signal to be reconstructed within the block, allowing for a reduction in training iterations.
4. During the initialization phase, use the neural network 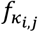 to fit the corresponding small block *m*_*i,j*_, with the objective function:

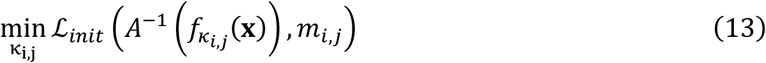

where *A*^−1^(·) is an operator that symmetrically crops the network’s output prediction volume from size (*D + R*) × (*F + R*) × (*F + R*) to size of *D* × *F* × *F*, for comparison with *m*_*i,j*_. The loss function is defined as follows:

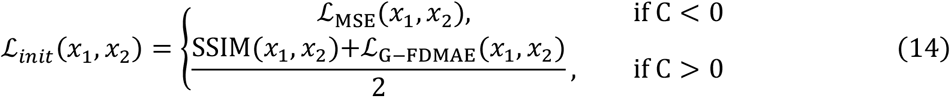

where *x*_1_ and *x*_2_ are two input images to be calculated.
5. During the reconstruction training phase, the neural network inherits the parameters *κ*_*i,j*_ obtained from the previous training phase, and integrates forward imaging process to solve the following equation:

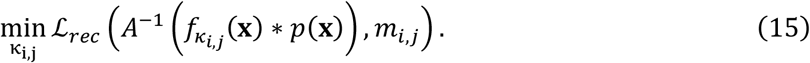

After training is completed, we can obtain the desired reconstruction result 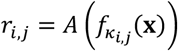.
6. In the stitching phase, the reconstruction results *r*_*i,j*_ from each block are stitched together to form the complete FOV. Since each block is reconstructed independently, regions closer to the center of each block have more complete information. Consequently, we have higher confidence in the reconstruction results for these regions compared to the edge regions. To account for this, we assign a smoothing mask *Q*_*i,j*_ to each block based on its position within the full FOV. The mask has a value of 1 in non-overlapping regions and decreases linearly with distance from the block center in overlapping regions. The final full-FOV result is computed as follows:

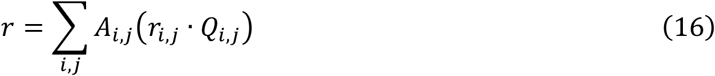

where *A*(·) is the operator that maps the *D* × *F* × *F* volume into the corresponding region of the full-FOV *D* × *W* × *H* volume.

